# Establishment of physiologically relevant oxygen gradients in microfluidic organ chips

**DOI:** 10.1101/2022.01.29.478323

**Authors:** Jennifer Grant, Elizabeth Lee, Micaela Almeida, Seongmin Kim, Nina LoGrande, Girija Goyal, Adama Marie Sesay, David T. Breault, Rachelle Prantil-Baun, Donald E. Ingber

## Abstract

*In vitro* models of human organs must accurately reconstitute oxygen concentrations and gradients that are observed *in vivo* to mimic gene expression, metabolism, and host-microbiome interactions. Here we describe a simple strategy to achieve physiologically relevant oxygen tension in a two-channel human small intestine-on-a-chip (Intestine Chip) lined with primary human duodenal epithelium and intestinal microvascular endothelium in parallel channels separated by a porous membrane while both channels are perfused with oxygenated medium. This strategy was developed using computer simulations that predicted lowering the oxygen permeability of poly-dimethlysiloxane (PDMS) chips in specified locations using a gas impermeable film will allow the cells to naturally decrease the oxygen concentration through aerobic respiration and reach steady-state oxygen levels < 36 mm Hg (< 5%) within the epithelial lumen. The approach was experimentally confirmed using chips with embedded oxygen sensors that maintained this stable oxygen gradient. Furthermore, Intestine Chips cultured with this approach supported formation of a villus epithelium interfaced with a continuous endothelium and maintained intestinal barrier integrity for 72 h. This strategy recapitulates *in vivo* functionality in an efficient, inexpensive, and scalable format that improves the robustness and translatability of Organ Chip technology for studies on microbiome as well as oxygen sensitivity.

## Introduction

Many living organs, such as the intestine, experience steep oxygen gradients that play a central role in health and disease, and this is particularly relevant for healthy host-microbiome interactions. The low-oxygen environment in the intestinal lumen and presence of a range of oxygen concentrations allows communities of complex anaerobic and aerobic microbes to thrive; however, the underlying human tissue requires higher levels of oxygen to survive. Most of what we know about the role of microbiome in medicine is based on genomic or metagenomic studies because it is extremely difficult to co-culture commensal bacteria in direct contact with human cells. In fact, oxygen is required for nearly every metabolic process in the human body, but vascular transport, molecular diffusion, and metabolism produce complex oxygen gradients in different organs that enable stable co-existence of complex communities of commensal microbes with living tissues. Recently, microfluidic organ-on-a-chip (Organ Chip) culture devices have been used to establish oxygen gradients that enable stable co-culture of human intestinal epithelium with a complex gut microbiome for days *in vitro*.^1,2^ However, these cultures required development of a specially fabricated hypoxia chamber and use of pumps and tubing that are not available in most laboratories, and this approach cannot be used with many Organ Chip operating systems. Thus, here we set out to explore whether a simpler approach can be developed to establish oxygen gradients within both lab fabricated and commercially available microfluidic Organ Chips *in vitro*.

The lumen or parenchyma of different organs operate within a characteristic oxygen concentration range. For example, the partial pressure of oxygen (pO_2_) ranges from 34-36 mm Hg (~5%) in the lumen of the small intestine^3^ and 3-11 mm Hg (0.4-1.5%) in the lumen of the large intestine^4^ to 34-54 mm Hg (4.5-7.1%) in the marrow compartment of bone.^5^ In all of these organs, nearby arterial and venous vessels experience higher oxygen concentrations of 70-100 mm Hg (9.2-13%) and 40 mm Hg (5.3%),^6^ respectively. Conversely, deviations from normal oxygen concentrations are characteristic of many disease pathologies, including diabetes, cancer, and ischemia.^7^ Thus, maintaining oxygen homeostasis is critical to maintain viability of different tissues and cells, as well as the commensal microbiome, which each have their own distinct oxygen requirements. Given the importance of oxygen status in the human body, it is therefore critical that *in vitro* organ and tissue models be cultured at physiologically relevant oxygen concentrations to ensure they optimally mimic *in vivo* functions.

Microfluidic culture systems can be precisely engineered to generate oxygen gradients through careful control over oxygen diffusion, convection, consumption, and generation. Perfused gas mixtures are commonly used to recreate physiologically relevant oxygen gradients in microfluidic culture systems. For example, we previously generated a hypoxia gradient in Intestine Chips, which contain two parallel microchannels separated by a porous membrane lined by intestinal epithelial and endothelial cells, by perfusing the lumen of the vascular channel with medium pre-exposed to a premixed gas containing 5% oxygen and culturing the chips in an anaerobic, nitrogen gas-filled atmosphere to which the epithelial channel had access via the gas permeable poly-dimethylsiloxane (PDMS) walls of the chip.^1,8,9^ A hypoxia gradient was also established in a two-channel Gut Chip lined with Caco-2 intestinal epithelial cells by increasing the thickness of the PDMS block above the cell-lined microchannel and perfusing anoxic medium through it while flowing oxygenated medium through a parallel channel separated from the first by a porous membrane.^10^ Continuous perfusion of anoxic and oxygenated media through adjacent culture chambers similarly has been used to generate oxygen gradients in the HuMiX Intestine Chip and GuMI platform.^2,11^ Although these methods recreated physiologically relevant hypoxia gradients, they are resource-intensive and prone to failure because gas tanks must be continuously monitored and replaced, gas connections often leak, the systems take hours to equilibrate, and the oxygen levels are difficult to control throughout the duration of the study. Additionally, routine laboratory procedures that are required for the experiment, such as changing media or moving the cultures to be viewed on a microscope, equilibrate the chips to aerobic conditions within minutes after removal from flow, which introduces variation into the experiments and greatly complicates the interpretation of results.

Alternative approaches that use aerobic respiration of cells cultured inside *in vitro* microsystems have the advantage that they do not require use of perfused gas mixtures. For example, a steady-state oxygen gradient can be established along the fluidic stream of a perfusion bioreactor when cells upstream in the bioreactor deplete oxygen for the cells downstream.^12,13^ However, this method generates an axial oxygen concentration gradient along the length of the fluidic channel, which does not permit analysis of key cell and tissue level functional parameters and biomarkers (e.g., epithelial barrier function, gene expression, cytokine release) because the cells along the fluidic stream are not maintained in a constant environment. Hypoxia gradients also have been established perpendicular to a cell layer in Transwell culture models by inserting an oxygen-impermeable plug over the apical side of the membrane and leaving the basal side open to atmospheric oxygen.^14,15^ But, these studies are limited because Transwells do not experience dynamic fluid flow or recapitulate physiological cell differentiation and tissue functions with the fidelity observed in Organ Chips.^16–18^

Here, we describe a simple strategy to achieve physiologically relevant oxygen tension in two-channel, microfluidic, Organ Chips by lowering the oxygen permeability in specific locations of the chip and allowing the cultured cells to further decrease the oxygen concentration through aerobic respiration. Our strategy generates a consistent oxygen gradient throughout the length of the chip when cultured in a conventional aerobic cell culture incubator and does not require anaerobic glove boxes or premixed gases. Furthermore, the oxygen levels remain stable when the chips are removed from the incubator. We used computational simulations of oxygen transport to determine that modifying commercially available PDMS Organ Chips by application of a polyvinylidene dichloride (PVDC) film along most of the surface of the chip allows the cells to naturally deplete oxygen in the microengineered lumen. We experimentally confirmed our approach using Intestine Chips with embedded oxygen sensors that quantify local oxygen concentrations. We verified the feasibility of this strategy for maintaining a hypoxic epithelial lumen by demonstrating no loss in intestinal barrier function over 72 h and maintenance of a viable villus epithelium interfaced with a continuous endothelium.

## Results and Discussion

### Simulating aerobic Intestine Chips

We previously described a two-channel, microfluidic, human Intestine Chip containing epithelial cells isolated from patient-derived organoids interfaced with human intestinal microvascular endothelium that recapitulates many anatomical and functional features of the *in vivo* small intestine (**Fig. 1a-c**).^16^ To explore potential methods to generate a stable oxygen gradient in this microfluidic Organ Chip, we first carried out computational simulations of the oxygen distribution. The Organ Chip we used for modeling is a commercially available (Emulate Inc.) PDMS microfluidic culture device (37.2 mm long x 16.2 mm wide) that contains parallel fluidic channels (15.46 mm long x 1 mm wide) separated by a porous PDMS membrane (50 μm thick; 7 μm pores). A rendering of the PDMS Organ Chip was imported into the simulation and the inlet and outlet regions were removed to increase computational efficiency (**Fig. 1b**). The pores in the membrane were also removed because the diffusion coefficient of oxygen in medium and PDMS are approximately the same (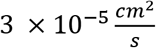 and 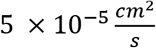, respectively)^10^, and removing the pores significantly decreased the geometric complexity of the model. The height of the epithelium was determined from morphometric analysis of cross-sections of Intestine Chips lined with primary human intestinal epithelium isolated from patient-derived organoids, as previously described.^1,19^ The endothelium was incorporated into the fluidic domain of the basal channel and not modeled as a separate geometry because it is very thin (<10 μm) compared to the rest of the chip. The simulation of oxygen distribution incorporated convection, diffusion, and the rate of endothelial and epithelial oxygen consumption. Air-saturated medium was flowed through the apical and basal channel at 60 μL/h, and the temperature, atmospheric pressure, diffusivity of oxygen in medium, and diffusivity of oxygen in PDMS, were set constant.

**Figure 1.**
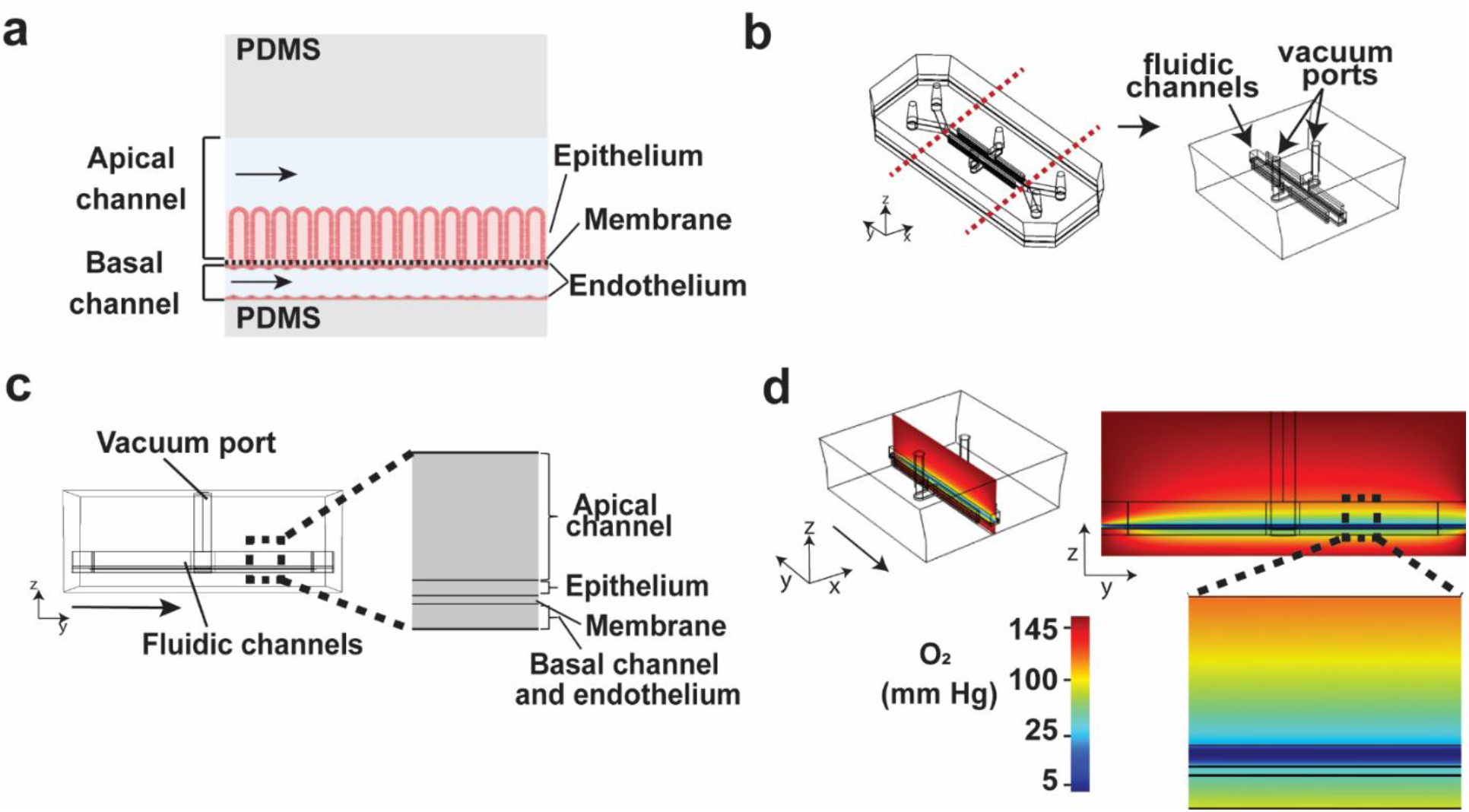
A) Side-view schematic of the two-channel Intestine Chip. Human intestinal epithelium is cultured on the porous membrane in the lumen (apical) microchannel and the bottom (basal) channel is lined with human vascular endothelium. Medium is perfused through the apical and basal channels. B) The Intestine Chip geometry simulated in COMSOL. The V-shaped inlet and outlet regions were removed to improve computational efficiency. The simulated geometry has two central fluidic channels that are separated by a porous membrane. The fluidic channels are sandwiched between two vacuum channels. C) Side-view image of the simulated Intestine Chip with the inset showing incorporation of the apical channel, epithelium, membrane, basal channel, and endothelium components in the model geometry. D) Steady-state heat map of oxygen distribution in the aerobic Intestine Chip. The oxygen concentration increases along the length of the apical channel away from the epithelium and the lumen does not sustain a physiologically relevant oxygen level.

The steady-state simulation of the aerobic Intestine Chip reveals that epithelial oxygen consumption generates a sharp oxygen gradient in both the apical and basal channel (**Fig. 1d**). The oxygen concentration rapidly increases across the height of the apical channel because the flux of oxygen through the PDMS exceeds the rate of epithelial oxygen consumption. To be relevant for the small intestine, the apical channel should sustain an oxygen concentration <35 mm Hg (<5%) above the epithelium.^3^ Therefore, additional modifications are required in order to achieve a physiologically relevant oxygen gradient in the Intestine Chip.

### Establishing Relevant Oxygen Gradients

As oxygen rapidly permeates through PDMS, we hypothesized that reducing the oxygen flux into the chip would allow the epithelium to naturally lower the oxygen concentration through aerobic respiration. We simulated coating the chip with a material that has low oxygen permeability (P_m_), while leaving the bottom of the basal channel open to atmospheric oxygen to generate a hypoxia gradient between the fluidic channels and lower the oxygen concentration above the epithelium in the upper channel. Specifically, we modeled coating the top surface of the chip with polyvinylidene chloride (PVDC) film because it is a flexible and self-adhering polymer film that has one of the lowest oxygen permeabilities at 100% humidity and 37 °C (**Fig. 2a**).^20,21^ Additionally, PVDC is transparent to allow for microscopy studies and is commercially manufactured as thin film (<0.025 mm) that is impenetrable to bacteria and mold,^22^ making it well-suited for use with Organ Chips. The steady-state simulation shows that coating the chip with PVDC lowers the oxygen concentration and that the chip sustains P_O2_<35 mm Hg (<~5% O2) within a 250 μm-tall region above the epithelium (**Fig. 2b**). Higher oxygen concentration is observed on the basal side of the chip due to the flux of oxygen through the uncoated region. This strategy generates a physiologically relevant hypoxia gradient perpendicular to cell layers cultured in the Intestine Chip by controlling the flux of oxygen into the device, while taking advantage of ongoing aerobic respiration in the cell layers to deplete oxygen. A sharp parabolic distribution of oxygen is observed at the inlet of the apical channel because the medium entering the chip is saturated with oxygen. The effect of oxygen consumption by the epithelium propagates into both the apical and basal channel; however, blocking the flux through the PDMS greatly lowers the oxygen concentration directly above the epithelium in the apical channel.

**Figure 2.**
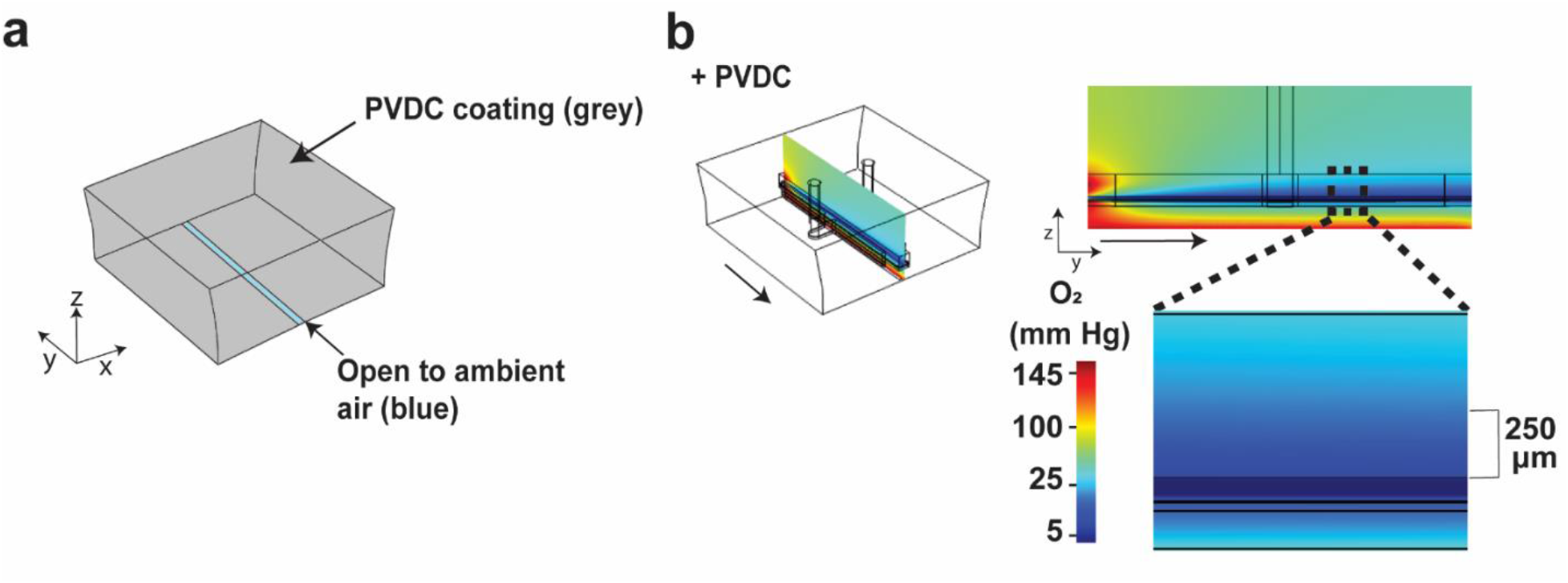
Reduction of the oxygen level and establishment of a physiologically relevant oxygen gradient in the Intestine Chip. A) To lower the oxygen level and establish an oxygen gradient, the chip is coated with PVDC film and the bottom of the basal channel is left open to ambient oxygen. B) Heat map of the steady-state oxygen concentration when the Intestine Chip is coated with PVDC film. The arrow represents the direction of medium flow.

We then simulated coating the chip with a polyethylene terephthalate (PET) film because PET has similar material properties to PVDC that also make it compatible with Organ Chip culture, however, PET is slightly more oxygen-permeable than PVDC. The steady-state simulation of the PET-coated chip shows that the oxygen concentration above the epithelium more rapidly increases than in the PVDC coated chip, and that the overall oxygen concentration in the PET coated chip is higher (**Suppl. Fig. S1**). Therefore, the oxygen permeability coefficient of the coating material affects the oxygen concentration in the apical channel and the steepness of the gradient between the apical and basal channel. Materials with lower oxygen permeability coefficients will therefore establish a steeper gradient with lower oxygen concentrations in the apical channel. The oxygen concentration can be tuned by selecting materials with different oxygen permeabilities and the coating material must be carefully selected to meet the oxygen requirements of the *in vitro* model. Together, the simulations indicate that a PVDC coating is the optimal choice for recapitulating the oxygen microenvironment of the intestine.

We also considered the impact of applying cyclic strain on the oxygen gradient, which is made possible in the Organ Chips we used by application of cyclic suction to hollow chambers that are located on either side of the cell-containing central channels (**Fig. 1b**). However, application of cyclic strain also requires that the vacuum channels parallel to the fluidic channels are exposed to the aerobic incubator atmosphere. Continuously stretching the membrane at 10% strain and 0.15 Hz lowers the P_O2_ in the vacuum channels by approximately 28% (P_O2_~ 100 mm Hg, 13%). The P_O2_ under cyclic strain was applied to all sides of the vacuum channel and the oxygen concentration was simulated at steady state (**Suppl. Fig. S2**). The simulation shows that cyclic strain eliminates the hypoxia gradient and significantly increases the oxygen concentration because the rate of oxygen entry into the fluidic channels from the vacuum channel exceeds the rate of aerobic respiration. Thus, using this method, a hypoxia gradient can only be generated under static conditions in these chips, however, this limitation could be overcome in the future by dip-coating the vacuum channels in PVDC, which would reduce the rate ambient air enters into the fluidic channels.^23^

### Experimentally Validating the Oxygen Gradients

To experimentally validate the anaerobic method defined by the computational simulations, we fabricated Organ Chips with embedded oxygen sensors that read out the *in-situ* oxygen concentrations at four distinct locations along the fluidic channels. The sensors are located at the top and the bottom of the apical and basal channels, and at two locations along their length (**Fig. 3a**). These sensor chips were not fabricated with vacuum channels because they interfered with the placement of the sensor spots. However, as previously described, the epithelium still undergoes spontaneous differentiation and expresses many *in vivo* features of the human small intestine under continuous fluidic flow without cyclic strain.^17^ We introduced human epithelium derived from intestinal organoids into the oxygen-sensing chips and cultured the chips for 19 days under continuous medium perfusion through both channels (60 μL/h) (**Suppl. Fig. S3**). The sensor chips were cultured with epithelium only because the endothelium has a minimal impact on the oxygen distribution in the Intestine Chip (**Suppl. Fig. S4**).

**Figure 3.**
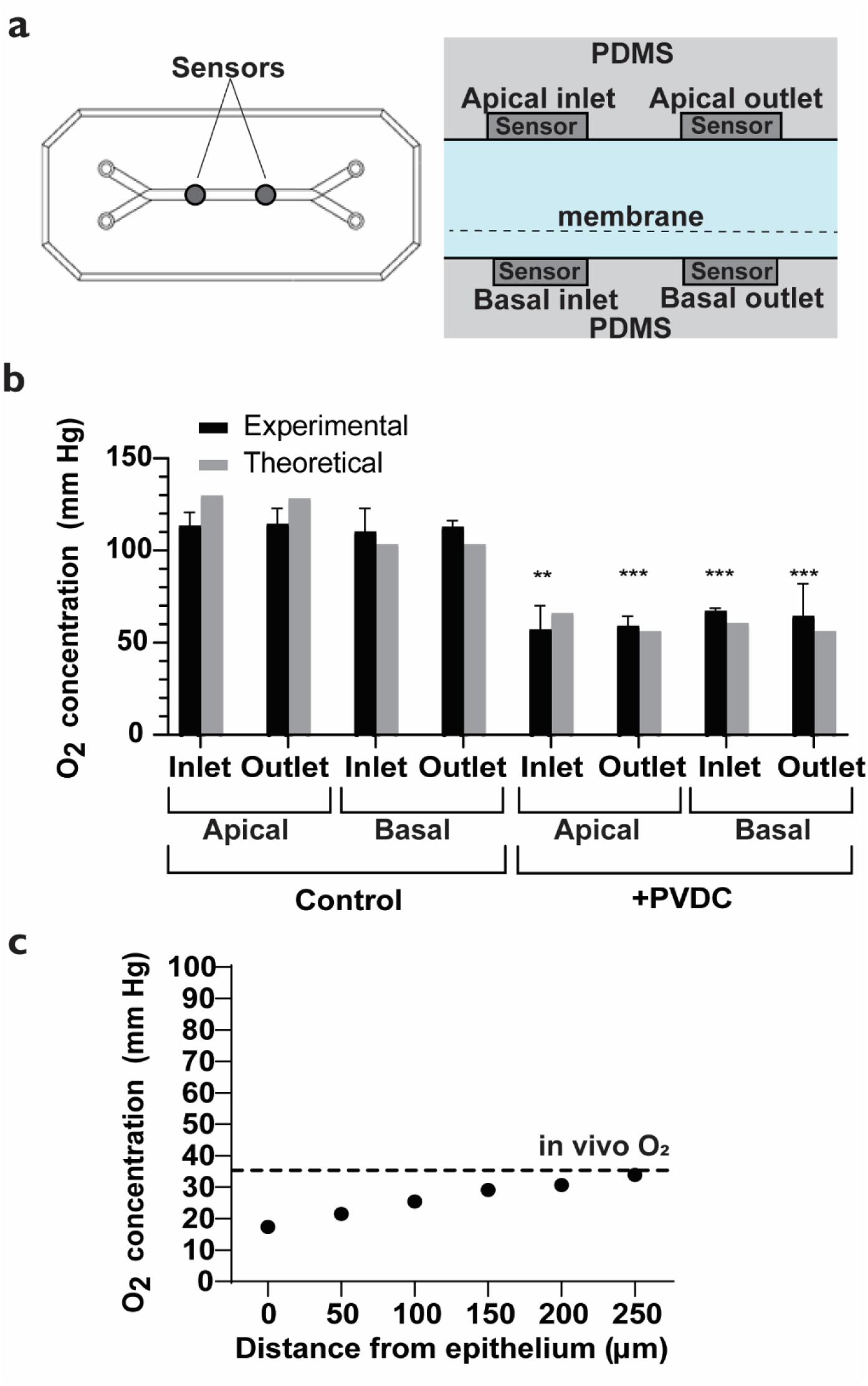
Experimentally validating the anaerobic strategy with oxygen-sensing Intestine Chips. A) A schematic representing the oxygen-sensing Intestine Chip with four oxygen sensor spots embedded in the top and the bottom of the apical channel at two locations in the middle of the chip. B) Oxygen concentration measured from the anaerobic and aerobic Intestine Chip. The experimentally measured oxygen is plotted alongside the theoretical oxygen concentration at the sensor locations (n=3 for experimental data). **, P<0.01; ***, P<0.001. C) A plot of the oxygen concentration at different distances away from the epithelium. The anaerobic Intestine Chip maintains a physiologically relevant oxygen concentration 250 μm above the epithelium.

We initiated anaerobic culture by wrapping the chips with PVDC film and introducing holes through this film along the bottom of the basal channel (**Suppl. Fig. S6**). The chips were placed in an aerobic cell culture incubator at 37°C and 5% CO_2_, and medium was flowed through both channels (60 μL/h) for 24 h. The oxygen concentration was measured from each sensor spot immediately after removal from flow (**Fig. 3b**). The PVDC-wrapped chips show a significant reduction in oxygen concentration at each sensor location, which validates our hypothesis that blocking oxygen flux into the chip lowers the oxygen concentration. We ran another simulation of the Intestine Chip that reflected the number of cells present in the chip as determined experimentally. In contrast to the previous steady-state simulations, we carried out this simulation to predict the oxygen concentration after removing the chip from flow for 2 min at 37°C because the oxygen sensor chips were removed from flow to acquire measurements and this procedure took approximately 2 minutes. The simulation analyzed the oxygen concentration at the same location of the sensor spots. We found that the simulation approximates the experimentally determined measurements within 90% accuracy at all sensor locations in both the aerobic and anaerobic chips. These findings confirm that the computational oxygen model is an effective tool to establish design criteria and better understand microfluidic culture systems.

The apical channel should sustain < 5% (< 35 mm Hg) oxygen to accurately recapitulate the microenvironment of the intestinal lumen^3^. As the oxygen sensors are placed on the top and bottom of the apical and basal channel, respectively, and do not represent the oxygen concentration directly above the epithelium, we analyzed this distribution computationally. A heat map of the steady-state oxygen distribution in the anaerobic Intestine Chip shows an oxygen gradient that decreases from the top of the apical channel to the epithelium (**Fig. 3c**). Plotting the simulated oxygen concentration at different distances away from the epithelium shows that the chip maintains oxygen < 35 mm Hg 0-250 μm above the epithelium, which is physiologically relevant for applications, such as bacterial co-culture and studies on oxygen sensitivity. This system establishes a hypoxia gradient across the epithelium and endothelium, which allows oxygenation of the endothelium while simultaneously providing an anaerobic environment above the epithelium for culture of both anaerobic and aerobic microbes.

This anaerobic strategy is inexpensive and uses laboratory equipment already inside tissue culture labs. Methods that use premixed gases require a large initial investment in capital equipment, demand more materials, and are difficult to scale up. These approaches have a large risk of experimental loss because they require continuously monitoring gas flows and connections, which can be prone to failure. Coating the Intestine Chip with a commercially produced film is a simple and scalable procedure that widens translatability of Organ Chip technology towards more oxygen-sensitive applications. Moreover, this approach is not limited to *in vitro* models of the intestine, as other systems such as the female genital tract, bone marrow, and colon experience *in vivo* oxygen gradients. Additionally, a variety of pathologies are associated with decreased oxygen levels or induced oxygen gradients, including inflammatory bowel syndrome^24^, cancer^25^, and reproductive diseases^26^, and thus, *in vitro* models that recapitulate the oxygen microenvironment should be able to more accurately represent diseased as well as healthy phenotypes.

Chips cultured with premixed gases experience wide fluctuations in oxygen concentration and take a long time to equilibrate back to steady-state. The ability to maintain steady oxygen levels outside of the incubator and without fluidic flow is important to carry out standard laboratory procedures (e.g., changing medium, microscopy imaging), and it enables experiments that demand extended time at room temperature. We removed the oxygen-sensing Intestine Chips from the incubator and measured a gradual increase in oxygen concentration over the course of 1 h (**Suppl. Fig. S5a**). The oxygen level increases because the rate of aerobic respiration in the intestine epithelium decreases by approximately 70% from 37°C to room temperature (22°C)^27^. We performed an additional computational simulation to predict the effect of removing the chips from flow and bringing them to room temperature, which incorporated this 70% reduction in aerobic respiration. The computational simulation predicted the experimental data with at least 90% accuracy at each time point, thus providing further validation of this simulation tool for assessing oxygen distributions on-chip (**Suppl. Fig. S5a**). The oxygen concentration sharply increases from the epithelial layer towards the top of the apical channel because the cells consume oxygen at a lower rate at room temperature (**Suppl. Fig. S5b**). Importantly, the region directly above the epithelium still maintains a consistent oxygen level over time after removal from flow (**Suppl. Fig. S5c**). The cells help maintain the oxygen level because they respond to perturbations in oxygen concentration. Because the cells obey Michaelis-Menten kinetics^28–30^, the PVDC-coated Intestine Chip will not experience wide fluctuations in oxygen concentrations over prolonged periods of time.

Our strategy for establishing a hypoxia gradient on-chip is more physiologically relevant than using premixed gases because aerobic respiration, diffusion, and convection, are the same mechanisms that generate hypoxia gradients *in vivo*. In the intestine, countercurrent blood flow in the villi and aerobic respiration by the epithelium and commensal microbiome together reduce P_O2_ in the lumen^31,32^. The flux of oxygen entering the lumen is offset by diffusive mixing, convective transport, and consumption of oxygen which contribute towards lowering the P_O2_. Similarly, our anaerobic Intestine Chip maintains low and stable oxygen levels by balancing cellular oxygen consumption with the flux of oxygen entering the chip to shift equilibrium to P_O2_ < 35 mm Hg (< 5%) above the epithelium.

### Maintenance of Intestinal Barrier Integrity

We then exposed Intestine Chips containing human small intestine epithelium and underlying vascular endothelium to the hypoxia gradient. Differential interference contrast (DIC) and immunofluorescence microscopic analysis confirmed that Intestine Chips cultured under hypoxia for 72 h maintained the villus epithelium interfaced with a continuous endothelium (**Fig. 4a, 4b**). We also evaluated the effect of hypoxia on intestinal barrier function by measuring changes in the apparent permeability coefficient (P_app_) and observed maintenance of barrier integrity for 72 h (**Fig. 4c**). Taken together, these data confirm that hypoxic Intestine Chips with primary human intestinal epithelium in contact with human endothelium maintain morphology and structural integrity.

**Figure 4.**
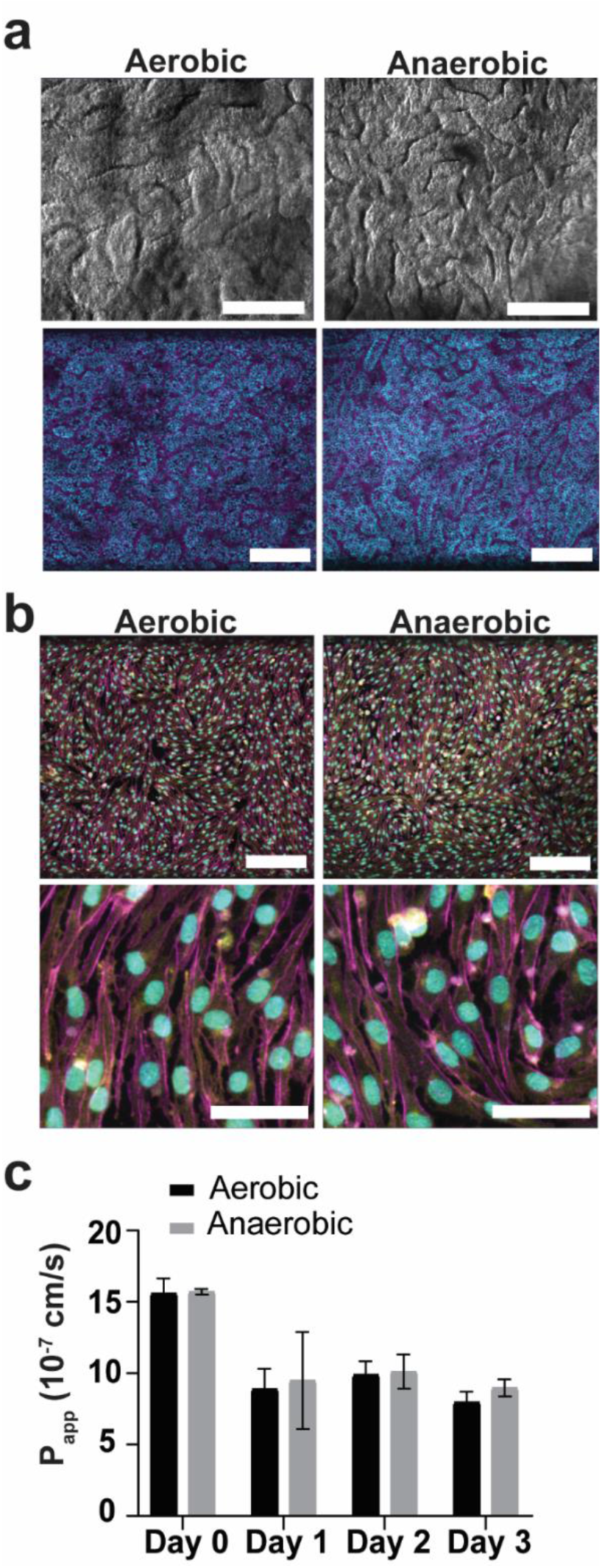
Culturing Intestine Chips under the hypoxia gradient. A) Top row: DIC microscopy views from above showing the villus morphology of the Intestine Chip after culturing under the hypoxia gradient for 72 h, compared to a chip cultured under an aerobic environment. Scale bar: 200 μm. Bottom: Immunofluorescence microscopy views from above showing the villus morphology of the Intestine Chip after culturing under anaerobic conditions for 72 h compared to aerobic conditions when stained for F-actin (magenta) and nuclei (blue). Scale bar: 200 μm. B) Immunofluorescence microscopy views showing maintenance of the endothelium layer when cultured under identical conditions as in A), when viewed from above and stained for VE-cadherin (yellow), nuclei (blue), and F-actin (magenta). Scale bar: 200 μm. Bottom: Scale bar: 50 μm. C) Intestinal barrier integrity measured for 72 h shows no change in apparent permeability (P_app_) between Intestine Chips cultured aerobically and anaerobically (n=3).

## Conclusion

We developed a simple strategy to achieve physiologically relevant oxygen tensions in human Intestine Chips by lowering the oxygen permeability in specific locations of the chip and allowing the cultured cells to decrease the oxygen concentration through aerobic respiration. Our approach is physiologically relevant because it uses respiration, convection, and diffusion to establish hypoxia, which are the same processes that drive the formation of oxygen gradients *in vivo*. This strategy is also scalable, cost-efficient, and easy to implement into standard laboratory workflows because an oxygen gradient can be generated in a conventional aerobic incubator and the chips can be removed from the incubator without significantly disrupting the oxygen distribution. This simplified approach for generating a hypoxia gradient in microfluidic Organ Chips enables the development of *in vitro* models that require a controllable oxygen environment, which could accelerate the discovery of therapeutic strategies for a broader range of human pathologies.

## Materials and Methods

### Computational Simulations

A SOLIDWORKS (Solidworks 2017) three-dimensional (3D) rendering of the microfluidic device was imported into COMSOL (COMSOL Multiphysics 5.5, COMSOL, Inc.) and the V-shaped inlet and outlet regions of the chip were removed to increase computation speed. The final simulated geometry was 15.46 mm long x 16.2 mm wide. Our Intestine Chip models also incorporate living intestinal epithelium and endothelium cultured on its top and bottom sides, respectively. The height of the epithelium (103 μm) and endothelium (< 10 μm) was experimentally determined from cross-sections of Intestine Chips. The endothelial layer is very thin compared to the membrane, epithelium, PDMS blocks, and fluidic channels, therefore, it was merged with the basal channel domain and not simulated as a separate geometry.

The 3D steady-state oxygen distribution was simulated by coupling laminar flow with dilute species transport. Fluid dynamics simulations were performed using the Navier-Stokes equations for incompressible flow and Fick’s second law was applied to model oxygen transport through the culture medium and PDMS.^10^ Oxygen consumption by the epithelium and endothelium was assumed to follow Michaelis-Menten kinetics, which incorporated the known oxygen consumption rate per epithelial and endothelial cell.^33–35^ The average number of epithelial and endothelial cells in the Intestine Chip were experimentally determined and incorporated into the maximum rate of oxygen consumption in the epithelium and endothelium lining the porous membrane, *V_max, epi_* and *V_max, endo_*, respectively. The effect of coating the outside of the chip with PVDC film was simulated by incorporating the oxygen flux through this material, which has a defined oxygen permeability and thickness.^36^ A custom meshing sequence generated finer mesh sizes at boundaries and corners. Steady-state simulations were performed under continuous fluid flow at 60 μL/h in the apical and basal channels and atmospheric pressure at 1 atm. Simulations in the cell culture incubator were performed at 37 °C and with 145 mm Hg atmospheric pO_2_. Time-dependent simulations of chips taken out of the cell culture incubator and brought to room temperature were performed with 0 μL/h fluidic flow, 1 atm atmospheric pressure, and 160 mm Hg atmospheric pO_2_. The steady-state solution from the simulation inside the cell culture incubator was used as the initial condition for the time-dependent simulation. Parameters and equations used in the simulations are provided in **Suppl. Tables S1** and **S2**, respectively.

### Cell Culture

Human small intestine (duodenal) epithelium was obtained and isolated from de-identified biopsy specimens obtained from Boston Children’s Hospital and all methods were in accordance with the Institutional Review Board of Boston Children’s Hospital (Protocol number IRB-P00000529). These specimens were collected from regions of the duodenum determined to be macroscopically healthy based on gross examination. From these biopsy samples, small intestine organoids were cultured and resuspended in Matrigel (356231, lot # 0090005; Corning) at approximately 1×10^6^ cells/mL in the presence of stem cell expansion medium. The stem cell expansion medium consisted of advanced Dulbecco’s modified Eagle medium F12 (12634-010; Thermo Fisher Scientific) supplemented with the following: L-WRN, Noggin, R-spondin conditioned medium (50% vol/vol^−1^), 1X GlutaMAX (35050-061; Thermo Fisher Scientific), 10 mM HEPES (15630-106; Thermo Fisher Scientific), 0.1 mg/mL primocin (antpm-1; InvivoGen), 1X B27 supplement (17504-044, Thermo Fisher Scientific), 1X N2 supplement (17502-048; Thermo Fisher Scientific), 10 mM nicotinamide (N0636; Sigma-Aldrich), 1.25 mM n-acetyl cysteine (A5099; Sigma-Aldrich), recombinant murine epidermal growth factor (50 ng/mL-1) (315-09; Peprotech), 10 nM human (Leu15)-gastrin I (G9145; Sigma-Aldrich), 500 nM A83-01 (2939; Tocris), and 10μM SB202190 (S7067; Sigma-Aldrich).

Intestine organoid cultures were maintained on a seven-day schedule with splitting or chip seeding occurring on the seventh day of growth. To isolate the organoids for splitting, Matrigel droplets containing the suspended organoids were incubated in cell recovery solution (354253; Corning) for 1 h on ice, then centrifuged at 300 x g for 5 min at 4 °C. The organoids were then resuspended in fresh Matrigel and mechanically fragmented with a pipette. The Matrigel-organoid suspension was plated in 45 *μ*L droplets on a coated 24-well plate (25-107; Olympus). Each droplet was covered with stem cell expansion medium supplemented with 10 μM Y-27632 (Y0503; Sigma-Aldrich). Stem cell expansion medium was replaced every 2 days.

The microfluidic Organ Chip devices were either purchased from Emulate, Inc (CHIP-S1 Stretchable Chip, RE00001024 Basic Research Kit; Emulate, Inc, Boston, MA) or fabricated in-house with oxygen sensing nanoparticles. To prepare the chips for cell culture, the surface of the fluidic channels and membrane were activated with ER1+ER2 (Emulate, Inc) in the presence of UV light, then coated with 200 μg/mL rat tail collagen type 1 (354236; Corning) and 1% reduced-growth factors Matrigel (356231, lot #0090005; Corning). The intestine organoids were isolated from Matrigel and resuspended in a chemical fragmentation buffer consisting of a 1:1 (vol:vol^−1^) solution of DPBS and TrypLE Express Enzyme (12605010; Thermo Fisher Scientific) supplemented with 10 μM Y-27632. The organoids were then enzymatically fragmented by placing them in a 37 °C water bath under agitation for 2 minutes. After the reaction was quenched with twice the volume of Defined Trypsin Inhibitor (R007100; Thermo Fisher), the cells were spun at 300 x g for 5 min at 4 °C and re-suspended in stem cell expansion medium supplemented with 10 μM Y-27632 to yield approximately 6 × 10^6^ cells/mL. The organoid fragments were seeded onto the apical channel of the Intestine Chip, and then incubated overnight at 37 °C under 5% CO_2_ to allow for cell adhesion to the upper surface of the ECM-coated porous membrane.

The next day, the chips were washed with stem cell expansion medium, inserted into Pod Portable Modules (RE00001024 Basic Research Kit; Emulate, Inc), then placed in a Zoë Culture Module (Emulate, Inc). Medium was flowed through each channel at 60 μL/hr and in some experiments cyclic strain was applied at 10% and 0.15 Hz. At day 15 of culture, stem cell expansion medium was replaced with differentiation medium consisting advanced Dulbecco’s modified Eagle medium F12 (12634-010; Thermo Fisher Scientific) supplemented with 1 *μ*g/mL Human recombinant R-spondin (120-38; Peprotech), 100 ng/mL Human recombinant Noggin (120-10C; Peprotech), 1X GlutaMAX, 10 mM HEPES,0.1 mg/mL primocin, 1X B27 supplement, 1X N2 supplement, 0.5 mM n-acetyl cysteine, 50 ng/mL recombinant murine epidermal growth factor, 10 nM human (Leu15)-gastrin I, 500 nM A83-01, and 10 μM DAPT (D5942; Sigma Aldrich).

Human intestinal microvascular endothelial cells (HIMECs) (ACBRI 666; Cell Systems) that were used to line the vascular channel of the chips were first expanded in microvascular endothelial cell growth medium (EGM™-2MV BulletKit, CC-3202; Lonza) in plastic tissue culture flasks (Falcon 353136 and 355001) until confluent. On the day of plating, the basal channels of the Intestine Chips were first rinsed with endothelial cell culture medium consisting of stem cell medium supplemented with components from EGM™-2 MV Microvascular Endothelial SingleQuots Kit (CC-4147; Lonza), and then 50 μL HIMECS (450,000 cells/chip) were introduced into the lower channel while placing the chip upside down and incubating at 37 °C for 1 hour. The basal channels were then washed with endothelial cell medium and the Intestine Chips were reinserted into the Pod Portable Modules and placed in the Zoë Culture Module (both from Emulate Inc.), which was set to flow differentiation medium through the apical channel (60 μL/hr) and endothelial cell medium through the basal channel (60 μL/hr).

### Anaerobic Chip Assembly and Culture

The anaerobic Intestine Chip was assembled by coating the chip with PVDC film, a material that has low oxygen permeability, while leaving the bottom side of the chip open to ambient air (Fig. S7). The chips were coated with PVDC film on day 20 of culture. Chips were detached from the Pod Portable Module, removed from the grey chip holder, and wrapped in two layers of polyvinylidene chloride (PVDC) film (Goodfellow USA #CV301150, 0.0125 mm thick). Excess film from the sides of the chip were removed with scissors and eight holes were punctured into the film along the bottom of the basal channel with a 1 mm diameter biopsy punch (Braintree Scientific #MTP-3331AA). The chip was placed back into the grey holder, ensuring that the film covered all sides and edges of the chip. Inlet and outlets were punctured into the film with needle-nose tweezers (McMaster Carr #5669A71), leaving the vacuum ports covered in PVDC film. A 1 inch strip of silicone film (McMaster Carr #7643A381) was placed on the bottom of the chip directly underneath the basal channel to allow ambient air to enter into the basal fluidic channel. The chips were perfused (60 μL/h) with HBSS with calcium and magnesium in the apical channel and expansion medium with endothelial components in the basal channel. Cyclic strain is not applied during anaerobic culture because it introduces ambient air that equilibrates with the medium and eliminates the hypoxia gradient (Fig. S2).

### Oxygen Sensor Chip Fabrication

The chip consists of 5 layers: an apical block, apical gasket, porous membrane, basal gasket, and basal block (Fig. S7). The membrane was produced by casting PDMS preset polymer (10:1 ratio, Sylgard 184) onto a photolithographically prepared master that has an array of 50 μm-tall posts and cured at 60 °C for 1 h in a drying oven. The apical block was made by casting PDMS (10:1 ratio) in an aluminum mold (7075 aluminum, DATRON neo), degassing for 1 h in a desiccator, and curing at 60 °C for one hour. The apical gasket and basal block were prepared by spin-coating PDMS (10:1) ratio on a polycarbonate sheet (McMaster Carr) to a thickness of 1 mm. Similarly, the basal gasket was prepared by spin-coating PDMS (10:1) to a thickness of 200 μm. The apical gasket, basal gasket, and basal block were degassed in a desiccator and cured at 60 °C for one hour. Excess PDMS from each component was removed with a plot cutter (Graphtec CE 5000-60). Oxygen sensors were integrated into the apical and basal PDMS blocks by pipetting 10 μL slurry of 10 mg/mL oxygen-sensing nanoparticles (OXNANO, Pyroscience) in chloroform (Sigma) into two spots embedded along the channel in each block. After the particles dried on the PDMS surface, another 10 μL of 10 mg/mL oxygen-sensing nanoparticles to ensure that the optical sensor is evenly spotted and that it completely covers the width of the main channel. The sensor spots were examined to confirm that there are no large air bubbles present. The chip with integrated oxygen sensors was assembled following a layer-by-layer approach. Each layer component was plasma treated for 45 seconds (Diener Plasma Sterilizer – Nano, Diener electronic GmbH + Co. KG Germany), aligned under a microscope, and bonded at 60 °C for one hour.

### Oxygen Measurements

The oxygen concentration was quantified in the oxygen sensing Intestine Chips using an optical fiber (SPFIB-BARE-CL2, PyroScience GmbH) connected to a FireSting-PRO oxygen meter (FSPRO-4, PyroScience GmbH) with an external temperature probe (Pt100, PyroScience GmbH). A manual background correction was performed on an Intestine Chip that did not contain oxygen sensor spots. A 2-point calibration was carried out with 0% and 100% oxygen standard solutions, where the 0% standard was obtained by degassing a 1 mg/mL slurry of oxygen-sensing nanoparticles (OXNANO, PyroScience GmbH) in water with argon. A background correction and 2-point calibration was performed before acquiring data from each time point and when switching between apical and basal measurements.

### Paracellular Permeability Measurements

50 μg/mL Cascade Blue hydrazide Trilithium Salt (550 Da) (C3239; Invitrogen) in HBSS was added to the apical channel of the Intestine Chip to measure barrier permeability. The tracer that diffused to the basal channel was measured in the effluents and calculated based on a logarithmic standard curve. Apparent paracellular permeability was calculated using the following equation:

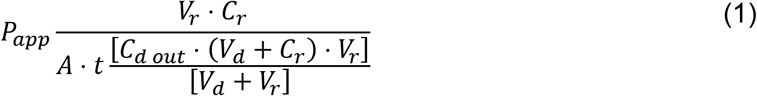

where *V_r_* is the volume of the receiving channel (basal channel effluent), *V_d_* is the volume of the dosing channel (apical channel), *C_r_* is the concentration of tracer in the receiving channel, *C_d out_* is the concentration of tracer in the dosing channel outlet (apical effluent), *A* is the area of the membrane (cm^2^), and *t* is time (sec).

### Morphological Studies

Intestine Chips were fixed with 4% paraformaldehyde (PFA) (RT157-4; Electron Microscopy Sciences) at room temperature 15 min 2x, then washed and stored in DPBS. The chips were either stained intact for whole-channel microscopy or sliced in 250 μm sections using a Vibratome (VT1000S; Leica). All samples were permeabilized with 0.2% Triton X-100 (T8787; Sigma) in DPBS for 30 min at room temperature under constant gravity flow, then blocked with 0.1% Triton X-100 and 10% Donkey Serum (D9663; Sigma) in DPBS statically for 1 hour at room temperature. The samples were then stained overnight at 4 °C with the following primary antibodies in an incubation and wash solution consisting of 1.5% bovine serum albumin (BSA) (A2153; Sigma) and 0.1% TritonX-100 in DPBS: Rabbit polyclonal to VE-cadherin/CD144 (ab33168, 1:100; Abcam) for intact chips and Rabbit polyclonal to Villin (PA5-17290, 1:100; Thermo Fisher) for sections. The next day the chips were washed 3 times with incubation and wash solution and left for 1 hour at room temperature with the secondary antibody, Donkey polyclonal to Rabbit IgG H&L, Alexa Fluor 647, (ab150075, 1:500, Abcam) and Phalloidin Alexa Fluor 488 (A12379, 1:200, Invitrogen) in the same solution. After the samples were washed 3 times with DPBS, they were stained with Hoechst 33342 (H3570, 1:2000; Life Technologies) for 30 min at room temperature. The samples were imaged using a Zeiss TIRF/ LSM 710 confocal microscope. Z-stacks were rendered and analyzed using Imaris.

### Statistical Analysis

Tests for statistically significant differences between groups were performed using a two-tailed t-test. Differences were considered significant when the P value was less than 0.05 (**, P<0.01; ***, P<0.001). All experimental results are expressed as means ± standard deviation (SD). Each experiment was conducted with a sample size of n=3 Intestine Chips per condition.

## ACKNOWLEDGMENTS

This work was supported by funding from FDA (75F40119C10098), Bill and Melinda Gates Foundation (OPP1173198), the Harvard Digestive Diseases Center (P30 DK034854) and the Wyss Institute for Biologically Inspired Engineering at Harvard University.

## POTENTIAL CONFLICTS

D.E.I. holds equity in Emulate Inc. and is a member of its board of directors and chairs its scientific advisory board.

