## Supplementary Information for "Establishment of physiologically relevant oxygen gradients in microfluidic organ chips"

**Suppl. Table S1.** Model parameters

| Parameter | Description | Value | Unit | Ref |
|---|---|---|---|---|
| $n_{epi}$ | Number of epithelial cells in the entire chip | $1 \times 10^6$ | *cells* | Experimentally determined |
| $n_{endo}$ | Number of endothelial cells in the entire chip | 60,000 | *cells* | Experimentally determined |
| $l$ | Length of the fluidic channels in the model | $15.46$ | $mm^2$ | |
| $f_{channel}$ | Fraction of total channel length represented by the model | $0.637$ | $mm$ | |
| $A_e$ | Cross-sectional surface area of epithelial cells | $1.03 \times 10^{-7}$ | $m^2$ | Experimentally determined |
| $r_{epi}$ | Oxygen consumption rate per cell | $3.5 \times 10^{-17}$ | $\frac{mol}{cell * s}$ | [1] |
| $r_{endo}$ | Maximum oxygen consumption of endothelial cells | $4 \times 10^{-17}$ | $\frac{mol}{cell * s}$ | [2] |
| $Q$ | Apical and basal inlet flow rates | $60$ | $\frac{\mu L}{h}$ | |
| $T$ | Incubator temperature | $37$ | $^\circ C$ | |
| $P$ | Atmospheric pressure in the incubator | $760$ | $mmHg$ | |
| $p_{O_2}$ | Partial pressure of oxygen in the incubator | $141$ | $mmHg$ | |
| $D_{medium}$ | Diffusion coefficient of oxygen in culture medium | $3 \times 10^{-5}$ | $\frac{cm^2}{s}$ | [3] |
| $D_{cell}$ | Diffusion coefficient of | $1 \times 10^{-6}$ | $\frac{cm^2}{s}$ | [4–7] |

| | | | | |
|---|---|---|---|---|
| | oxygen in epithelium | | | |
| $D_{pdms}$ | Diffusion coefficient of oxygen in PDMS | $5 \times 10^{-5}$ | $\frac{cm^2}{s}$ | [3] |
| $cO_{2sat}$ | Oxygen concentration at saturation in culture medium | $0.2097$ | $\frac{mol}{m^3}$ | |
| $K_{m,epi}$ | Concentration of oxygen at half maximum rate of epithelial oxygen consumption | $0.001$ | $\frac{mol}{m^3}$ | [8] |
| $K_{m,endo}$ | Concentration of oxygen at half maximum rate of endothelial oxygen consumption | .0007 | $\frac{mmol}{L}$ | [2] |
| $P_{PVDC}$ | Oxygen permeability coefficient of PVDC | $5.1 \times 10^{-18}$ | $\frac{mol * m}{m^2 * s * Pa}$ | [9] |
| $P_{PET}$ | Oxygen permeability coefficient of oriented PET | $7.6 \times 10^{-17}$ | $\frac{mol * m}{m^2 * s * Pa}$ | [10] |
| $h_{PVDC}$ | Thickness of PVDC film | $0.025$ | $mm$ | |
| $k_{O_2}$ | Henry's coefficient for oxygen | $9.901 \times 10^{-6}$ | $\frac{mol}{m^3 * Pa}$ | [11] |
| $v_b$ | Volume of the basal channel | 3.093 | $mm^3$ | |

**Suppl. Table S2**. Model equations

| Description | Function | Unit | Ref |
|---|---|---|---|
| Number of epithelial cells represented in the model | $n_{epi\_cells,model} = f_{channel} * n_{epi}$ | *cells* | |
| Number of endothelial cells represented in the model | $n_{endo\_cells,model} = f_{channel} * n_{epi}$ | *cells* | |
| Volume of the epithelium | $v_a = A_e * l$ | $m^3$ | |
| Density of epithelial cells | $\rho_{epi} = \frac{n_{epi\_cells,model}}{v_a}$ | $\frac{cells}{m^3}$ | |
| Density of endothelial cells | $\rho_{endo} = \frac{n_{endo\_cells,model}}{v_b}$ | $\frac{cells}{m^3}$ | |
| Maximum oxygen consumption rate of the epithelium | $V_{max,epi} = r_{epi} * \rho_{epi}$ | $\frac{mol}{m^3 * s}$ | |
| Maximum oxygen consumption rate of the endothelium | $V_{max,endo} = r_{endo} * \rho_{endo}$ | $\frac{mol}{m^3 * s}$ | |
| Rate of epithelial oxygen consumption | $N_{epi} = \frac{V_{max,epi} * c_{O2}}{K_{m,epi} + c_{O2}}$ | $\frac{mol}{m^3 * s}$ | |
| Rate of endothelial oxygen consumption | $N_{endo} = \frac{V_{max,endo} * c_{O2}}{K_{m,endo} + c_{O2}}$ | $\frac{mol}{m^3 * s}$ | |
| Global mass transfer coefficient | $K_{0_2,PVDC} = \frac{P_{PVDC}}{h_{PVDC}}$ | $\frac{s * mol}{kg * m}$ | 12 |
| Flux of oxygen through PVDC film | $N_{O_{2,PVDC}} = K_{0_2,PVDC} \left(p_{O_2} - k_{O_2} * c_{O_2}\right)$ | $\frac{mol}{m^2 * s}$ | 12 |

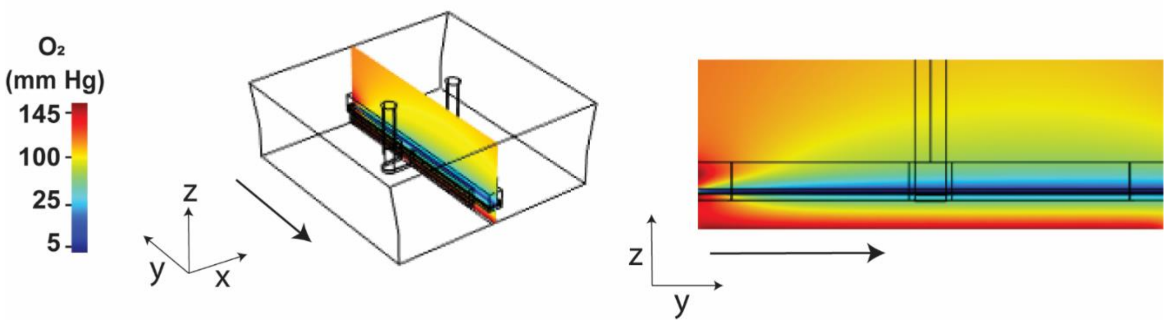


**Suppl. Fig. S1.** Steady-state oxygen concentration of the Intestine Chip coated in PET film.

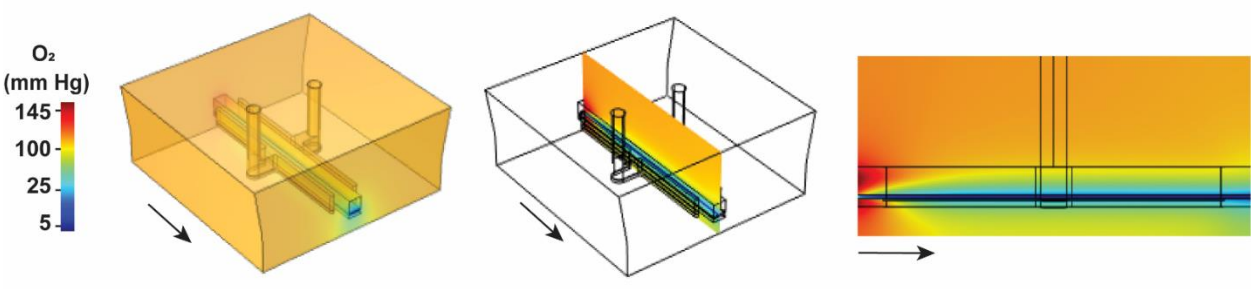


**Suppl. Fig. S2.** Steady-state solution of Intestine Chip when cyclic strain is applied. Cyclic strain introduces ambient air into the vacuum channels, which equilibrate with the medium and eliminate the hypoxia gradient. The arrows represent the direction of medium flow.

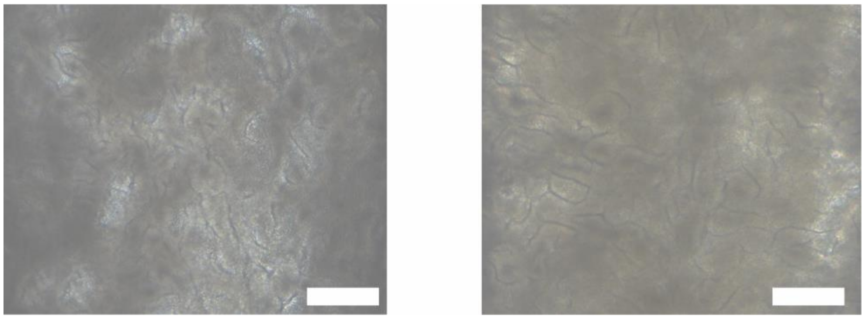

**Suppl. Fig. S3.** Epithelium on oxygen-sensing chip (left) and on standard chip without oxygen sensors (right). Scale bar: 200 µm.

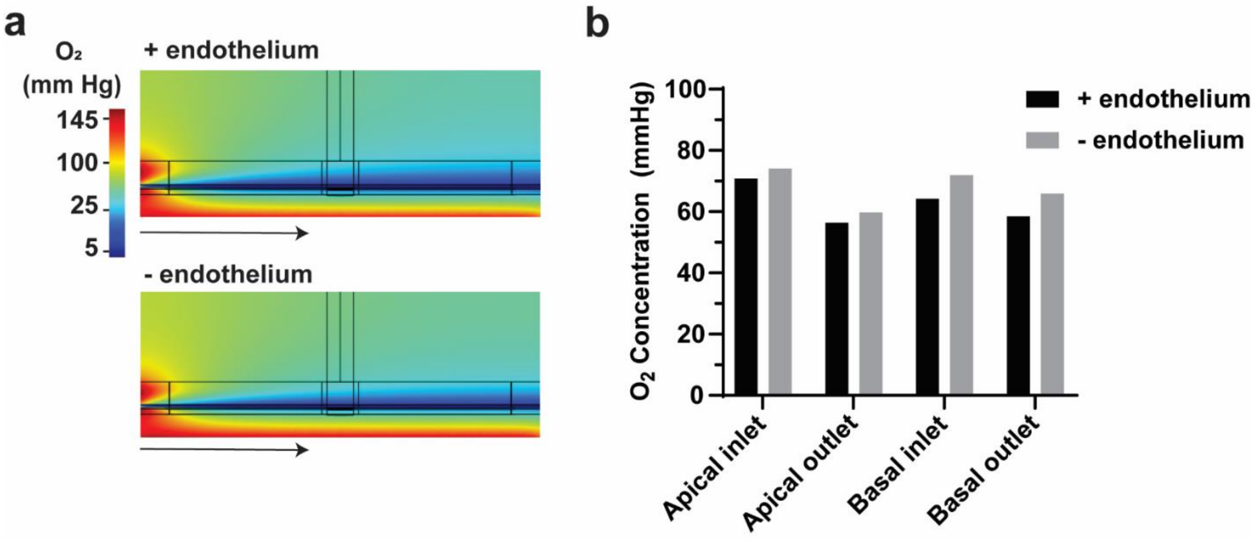


**Suppl. Fig. S4.** The endothelium has minimal impact on oxygen distribution and concentration. A) Steady-state oxygen distribution in Intestine Chips with (top) and without (bottom) endothelium. B) Plotting the theoretical oxygen concentration at four locations on the Intestine Chip with (black) and without (grey) endothelium.

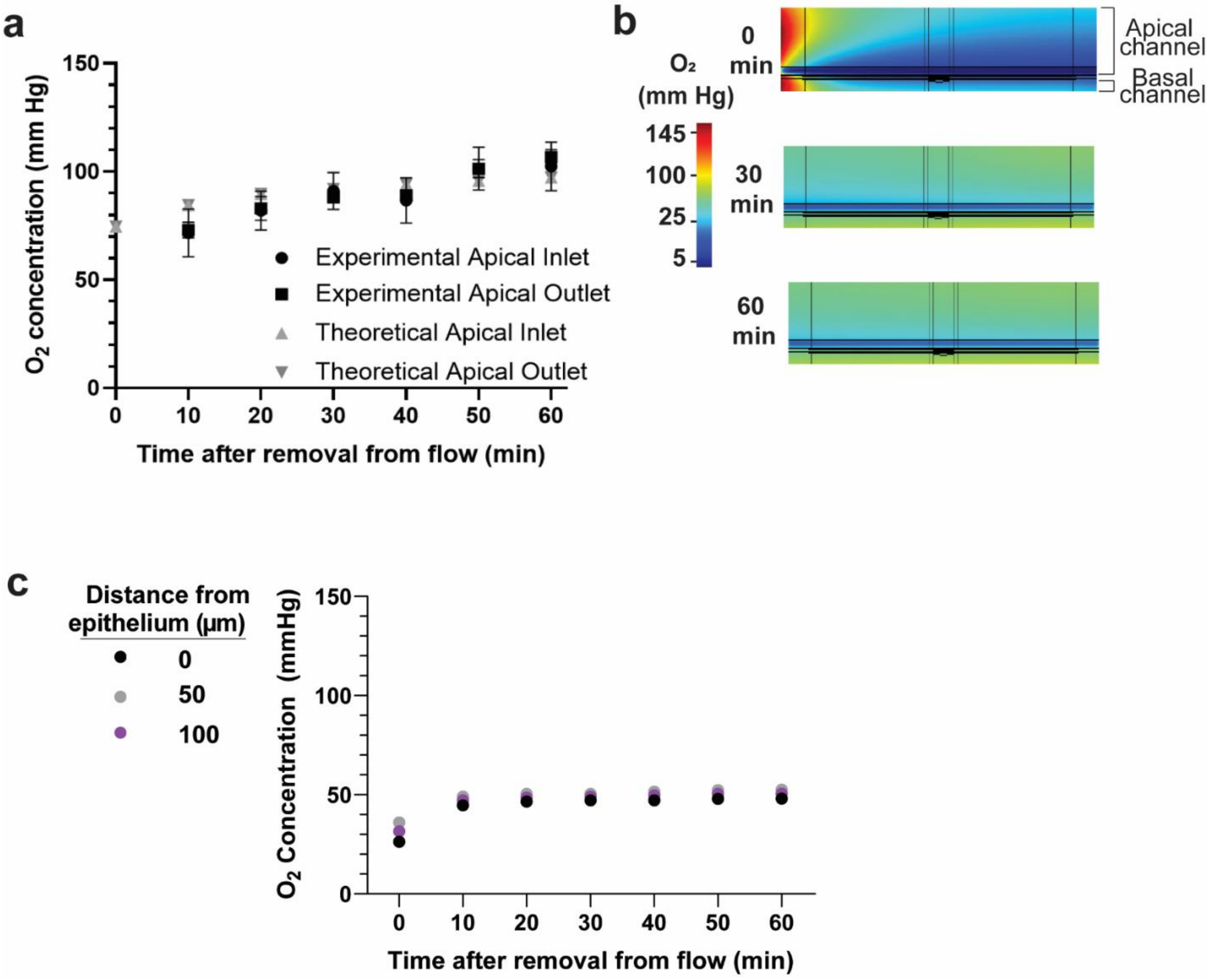


**Suppl. Fig. S5.** Effect of removing the chips from flow and bringing to room temperature. A) Experimentally measuring oxygen after chips are removed from flow and brought to room temperature. B) COMSOL simulations of removing the chips from flow and bringing them to room temperature. There is a gradient of oxygen concentration in the apical channel. Lower oxygen levels are closer to the epithelium. C) Theoretical measurements from inside the apical channel show that the chip is able to maintain stable oxygen levels 60 min after removal from flow.

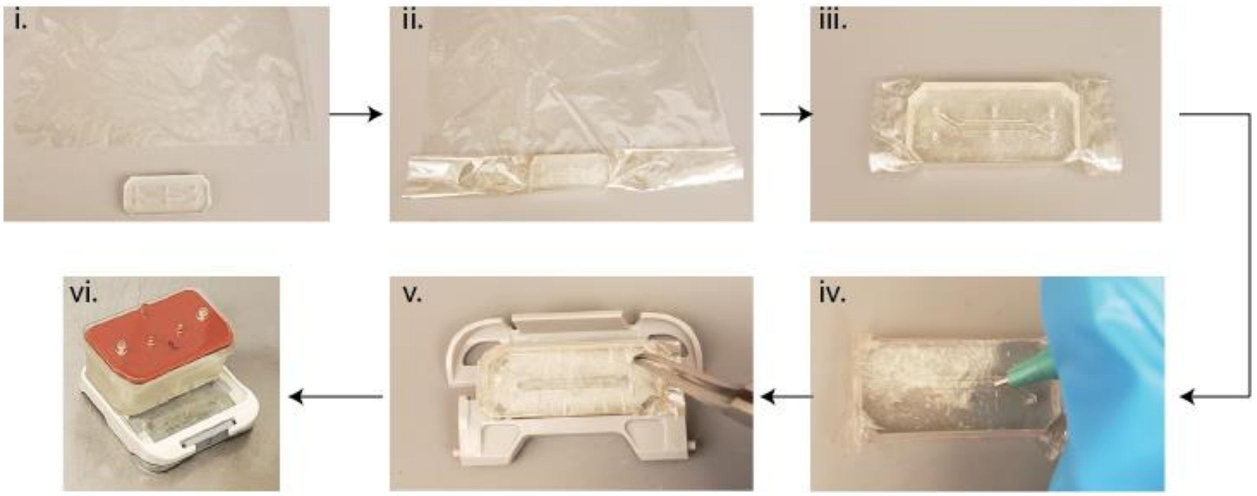


**Suppl. Fig. S6.** Procedure for coating the chip with PVDC film. i) cut a square of PVDC film, ii) tightly wrap the Intestine Chip with two layers of film, iii) trim excess film from the edges, iv) puncture 8 equidistant holes along the bottom of the basal channel with a 1 mm biopsy punch, v) introduce holes into the inlets and outlets, iv) insert the chip into the Pod.

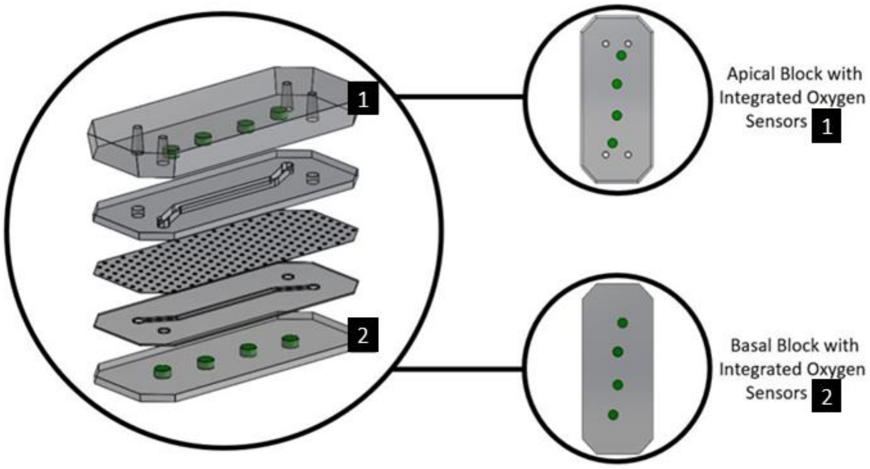


**Suppl. Fig. S7.** Design of the fabricated oxygen-sensing chip. The apical and basal blocks have multiple embedded oxygen sensors that face the main fluidic channels of the chip.
